# Super-Resolution Ultrasound Bubble Tracking for Preclinical and Clinical Multiparametric Tumor Characterization

**DOI:** 10.1101/203935

**Authors:** Tatjana Opacic, Stefanie Dencks, Benjamin Theek, Marion Piepenbrock, Dimitri Ackermann, Anne Rix, Twan Lammers, Elmar Stickeler, Stefan Delorme, Georg Schmitz, Fabian Kiessling

## Abstract

Super-resolution imaging methods promote tissue characterization beyond the spatial resolution limits of the devices and bridge the gap between histopathological analysis and non-invasive imaging. Here, we introduce Ultrasound Bubble Tracking (UBT) as an easily applicable and robust new tool to morphologically and functionally characterize fine vascular networks in tumors at super-resolution. In tumor-bearing mice and for the first time in patients, we demonstrate that within less than one minute scan time UBT can be realized using conventional preclinical and clinical ultrasound devices. In this context, next to highly detailed images of tumor microvascularization and the reliable quantification of relative blood volume and perfusion, UBT provides access to multiple new functional and morphological parameters that showed superior performance in discriminating tumors with different vascular phenotypes. Furthermore, our initial clinical results indicate that UBT is a highly translational technology with strong potential for the multiparametric characterization of tumors and the assessment of therapy response.

## Introduction

Ultrasound (US) is among the most frequently used diagnostic modalities in clinical routine, and its spatial and temporal resolution as well as tissue contrast have been steadily improved. The application of gas-filled microbubbles (MB) as US contrast agents further enhances the diagnostic accuracy of US by adding morphological and functional information about the tissue vascularization ^1^. This is particularly relevant in oncology, since the vascular structure of tumors contains essential information for their differential diagnosis ^2–4^, prognostication ^5^, and for the prediction and monitoring of therapy responses ^6–8^. In particular, some vascular features have already been shown to be capable of identifying patients not responding to antiangiogenic therapy ^9^, who, then, can be reoriented towards alternative approaches ^10^.

Different qualitative and quantitative techniques have been developed to extract the information about tumor vasculature contained in contrast-enhanced US (CEUS) scans. However, in state-of-the-art CEUS imaging, e.g. using Maximum Intensity Over Time (MIOT) ^11^ or replenishment kinetics analysis ^12^, voxels are usually much larger than the majority of tumor blood vessels, whose diameters are in the range of 5-80 μm ^13^. This limitation in the spatial resolution makes it difficult to gain a comprehensive overview of the vascular architecture and its heterogeneity. In addition, since the probability is high that every voxel contains at least one blood vessel, the tumor vascularization tends to be overestimated whenever the relative blood volume (rBV) is determined based on the area that exhibits MB signals ^14^. Voxel-wise analyses are further complicated by high background noise, which can make the assessment of functional vascular parameters difficult and unreliable at the single voxel level ^15^.

To overcome these issues, several postprocessing algorithms for CEUS image analysis have recently been proposed to reveal and quantify vascular features at super-resolution, which means at a resolution beyond the resolution limits of the device ^16,17^. Here, individual MB are localized, and a line with the thickness of a MB is drawn connecting the most closely localized MB in two subsequent frames. This line represents the track of a MB and thus, the course of a (micro) vessel. The approach was successfully applied to characterize MB flow tracks in brain ^16^ and ear vessels ^17^. However, in case of ambiguous assignment possibilities, this approach could lead to underestimation of flow velocities and might be particularly difficult to apply to more complex tumor vascular networks. Therefore, Errico and colleagues ^16^ used an experimental imaging system with a very high frame rate (500 frames per second) to avoid ambiguous assignments. However, comparable frame rates are not realized in the majority of clinical US systems so far, which makes clinical translation of this method difficult.

Therefore, we present here an alternative super-resolution CEUS approach called “Ultrasound Bubble Tracking (UBT)”, which is an advanced tracking technique that is adapted to clinical settings. With UBT, within less than a minute and using a conventional US devices operating at standard frame rates, super-resolution images and novel parameters could be extracted, which enabled an accurate discrimination of tumors with different vascular phenotypes. Furthermore, the preliminary clinical data that are presented in this manuscript show that rapid translation of UBT is realistic and that this technology may improve the diagnostic potential of CEUS in future clinical practice.

## Results

### Ultrasound Bubble Tracking for structural imaging of tumor vasculature

The UBT method reliably captured the movement of MB in tumors and could be successfully applied to all contrast-enhanced scans using a commercial US device operating at frame rates of approximately 50 frames/s and only required measurement times of 40 s (for more information about the algorithm, see Fig. 1a. as well as the detailed description in the “Materials and Methods” section). A spatial resolution of approximately 5 μm was achieved, and the vascular architecture of tumors was visualized in fine detail (Fig. 1b). Functional parameters were calculated for single vessels and combined with textural features, which so far could not been obtained with standard CEUS imaging (Fig. 1c).

**Fig. 1.**
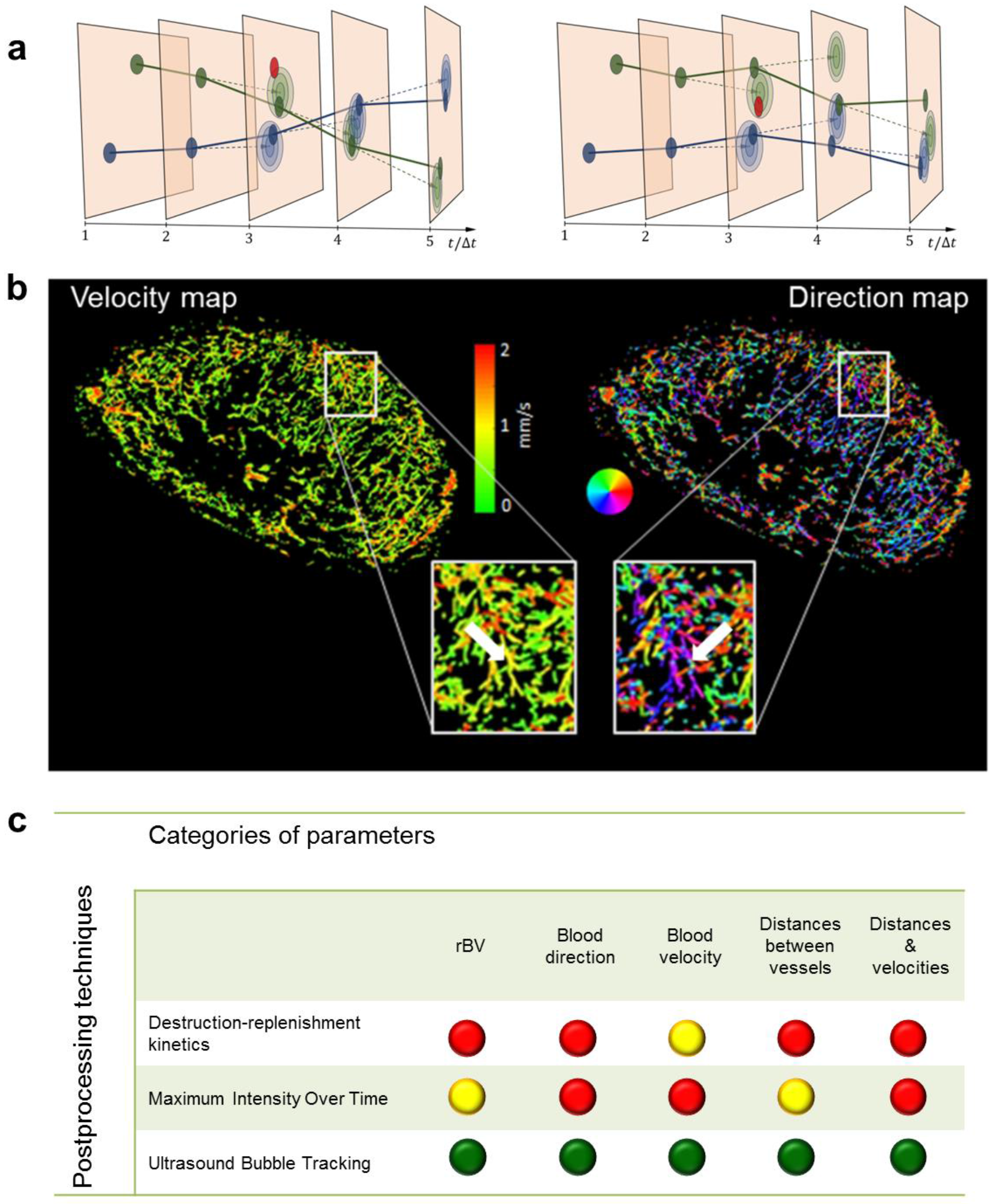
Ultrasound Bubble Tracking: Principle, examples and assessable parameters. (a) Sketch illustrating the principle of UBT. The filled circles mark the positions of detected MB. The red circles indicate detected MB supposed to be false alarms. The colors (blue/green) indicate the association of the MB to different tracks. One possible association of MB tracks is shown in the left diagram, another one in the right. The lighter ellipses indicate the probability density functions for the positions predicted by a linear motion model. From these, the likelihoods of the detected positions for an association are determined. The Markov Chain Monte Carlo Data Association (MCMCDA) algorithm searches for the association that maximizes the posterior probability. This also accounts for prior probabilities, like e.g. the probability of false alarms. Taking these analyses into account, in this example, the left association is more probable than the right one. (b) Super-resolution ultrasound images of an A431 tumor provide detailed information on the microvascular architecture including insights into vascular connectivity and the number of vascular branching points (see arrows in magnifications). Functional information such as MB velocities (left image) and MB flow directions (right image; color-coding illustrating the direction of flow according to the colored circle) can be determined for each individual vessel and evaluated together with the morphological characteristics. (c) Overview of the parameter classes obtained with UBT and their accessibility with standard contrast-enhanced ultrasound methods (green dot: quantitative and robust assessment of a parameter is possible; yellow dot: the information is available but its assessment is less robust, less accurate or not quantitative; red dot: the parameter cannot be obtained with the respective method).

At super-resolution, differences in the vascular texture in different tumor models could be clearly depicted. For instance, in line with histological staining, the highly angiogenic A431 tumors showed a fine network of very small vessels, homogeneously distributed throughout the entire tumor tissue (Fig. 1b, 2). In contrast, A549 tumors, which are known to be less vascularized and characterized by a more mature vascular system, displayed a higher vascular hierarchy in super-resolution UBT images, with larger vessels at the periphery, that branch into smaller vessels towards the tumor center. MLS tumors with their heterogeneous vascular pattern were most difficult to classify. In the super-resolution UBT images, as in histology, they were characterized by highly and poorly vascularized regions, and by more or less dense and branched vascular areas (Fig. 2).

**Fig. 2.**
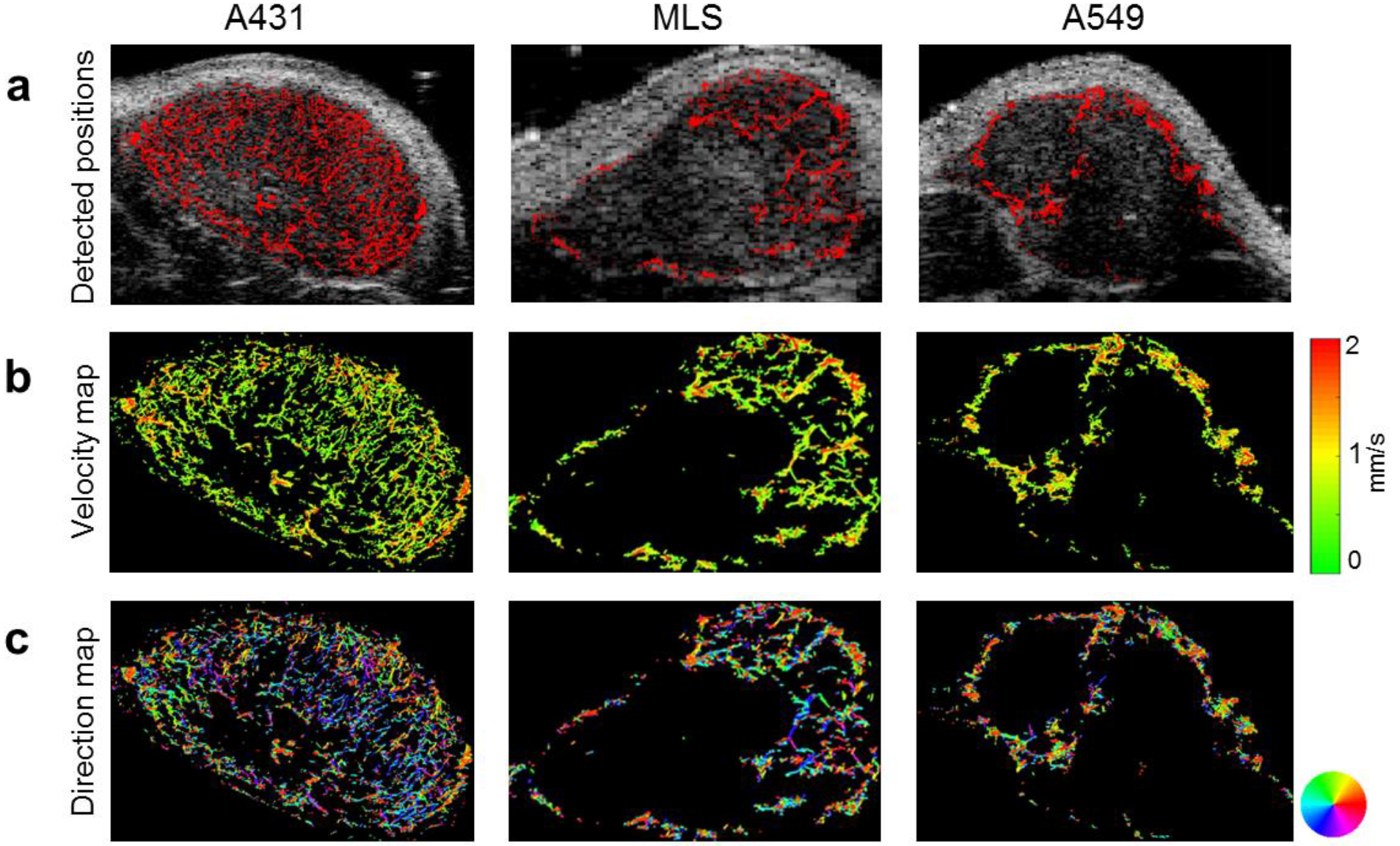
UBT-based parameter maps of tumors with different vascular phenotypes. The color-coded maps indicate the detected positions of MB overlaid on the B-mode images, representing the relative blood volume (a), individual MB velocities (b) and directions of MB flow (c). The three tumor models can be distinguished based on their different vascular patterns and quantitative textural analysis can be performed based on the super-resolution parameter maps.

As an additional feature of the UBT approach, velocities of MB and their directions of movement can be calculated for individual vessels and displayed in parametric maps (Fig. 1b). In these parametric direction maps information about arterial and venous supply as well as branching and connections between vessels is provided. While the velocity profiles of the tumors were similar, the analysis of flow directions showed differences. The parametric direction maps of A431 tumors indicated that blood flow in the central fine vascular network was predominantly directed towards the tumor core, while in the periphery the blood flow directions were chaotic (Fig. 1b). Thus, a balance between feeding and draining vessels was found in the periphery, while draining vessels were less apparent in the center. This lack of venous drainage is known to be a typical characteristic of highly angiogenic tumors ^18^. In contrary, the parametric maps of A549 and MLS tumors, which have a more mature vascularization (Supplementary Fig. 1), showed a balanced mixture of feeding and draining vessels (Fig. 2c), which is also reflected by their higher *flow direction entropy* values (Fig. 3b).

**Fig. 3.**
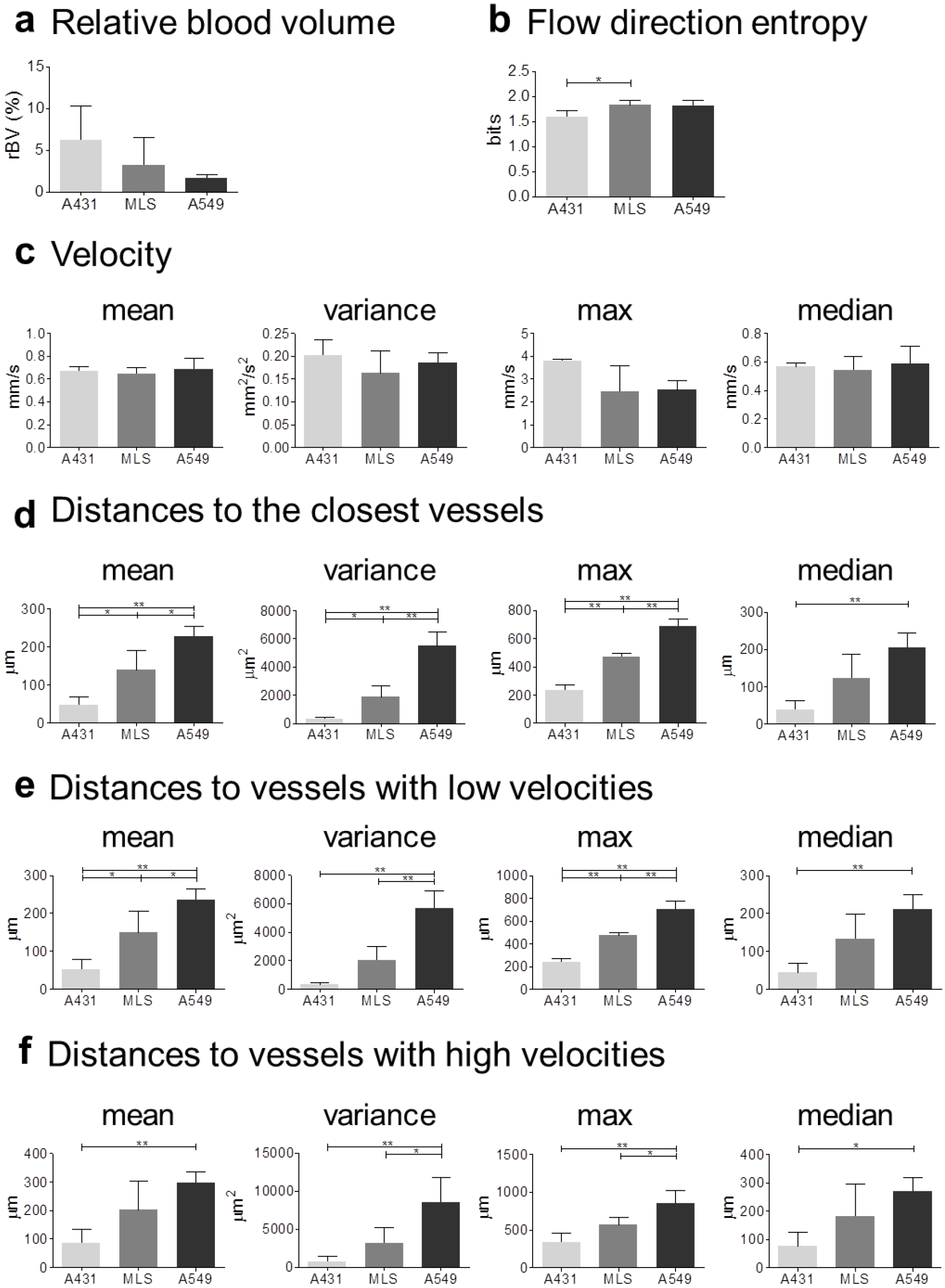
Comparison of UBT parameters. While *rBV* and MB *velocities* did not differ significantly between A431, MLS and A549 tumors (a and c), the tumor models could be distinguished using the parameters of *flow direction entropy*, *distances to the closest vessel*, and the parameters that combined velocity and distance information i.e *distances to vessels with low and high velocities* (b, d, e, and f). Only parameters that could distinguish all three tumor models were used for further analysis. For all bar plots shown, data are expressed as the mean ± s.d. (**=p<0.01; *=p<0.05; by one-way ANOVA with Bonferroni post-hoc analysis).

### Characterization of vascular tumor phenotypes using Ultrasound Bubble Tracking

While some UBT parameters can also be obtained by state-of-the-art CEUS postprocessing techniques, others represent new parameter classes that so far have been difficult to assess (Fig. 1c and Supplementary Table 1). Parameters determined by UBT include the *relative blood volume (rBV)*, the mean, variance, maximum and median values of MB *velocities*, *distances to the closest vessel*, and *distances to vessels with low and high velocities*, as well as the *flow direction entropy* as a measure for the organization of the vessel networks.

In good agreement with their histological vascular phenotypes, A431 tumors had the highest *rBV* values, followed by MLS and A549 tumors (Fig. 3a). However, due to the high heterogeneity of *rBV* within the respective tumor models, our group sizes were not sufficiently large to generate significant differences.

As expected from the visual inspection of the parametric maps (Fig 2c), the *flow direction entropy* values, describing the order of blood direction profiles, were lower in A431 than in MLS and A549 tumors. While this parameter could not unambiguously separate all groups, significant differences were found between A431 and MLS tumors (Fig. 3b).

Surprisingly, parameters related to MB *velocity* were very similar across the tumor models, indicating that, despite their different angiogenic phenotypes, these tumors tend to preserve a similar flow pattern (Fig. 3c).

The textural parameters of the tumor vascularization showed a significantly higher discriminatory potential (Fig. 3d-f). In this context, super-resolution images obtained by UBT enabled us to determine the *distances to the closest vessel* and to evaluate their mean, variance, maximum and median values. Strikingly, the first three of the above parameters had the power to discriminate all three tumor groups. However, among all distance parameters, the maximum of *distances to the closest vessel* was one of the best performing ones, which precisely discriminated all three tumor models (A431 vs. MLS, p< 0.01; A431 vs. A549, p< 0.001 and A549 vs. MLS, p<0.01) (Fig. 3d).

Two new parameter classes were introduced, which combine textural and functional information, i.e. 1) *distances to vessels with low velocities* and 2) *distances to vessels with high velocities*. While based on the parameters associated with *distances to vessels with high velocities* only one or two out of three possible combinations revealed significant differences, mean and maximum values of *distances to vessels with low velocities* differed significantly between all tumor models (Fig. 3e,f). Thus, the latter parameters were considered in the further analysis.

### Confusion and correlation matrices of novel parameters obtained by UBT

Parameters that could distinguish all three tumor models by statistical evaluation with one-way ANOVA and Bonferroni post-hoc tests were used for further analyses. These parameters were: 1) mean, 2) variance and 3) maximum of *distances to the closest vessel* as well as 4) mean and 5) maximum of *distances to vessels with low velocities* (Fig. 4a).

**Fig. 4.**
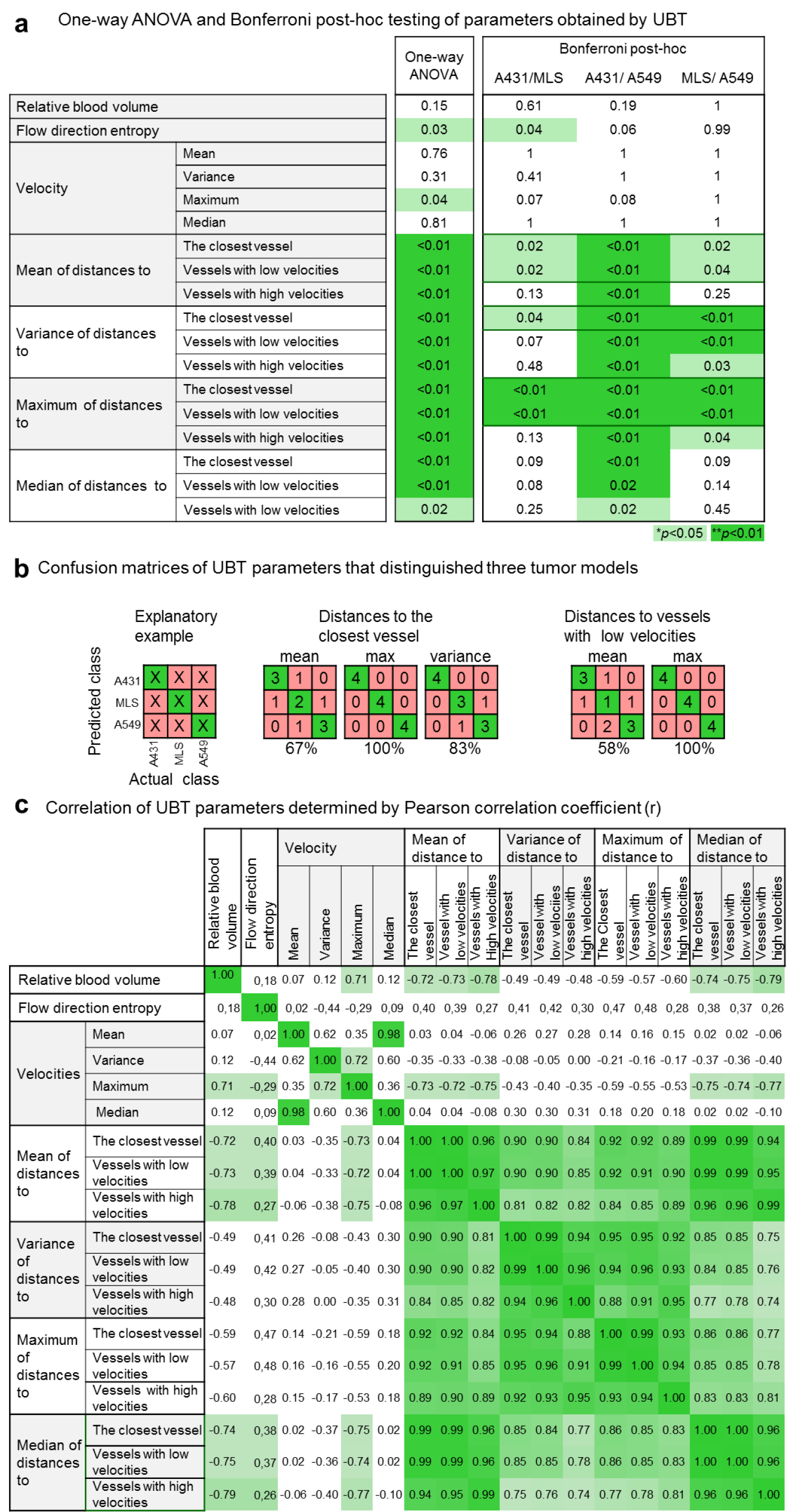
Capability of UBT parameters to distinguish tumors with different vascular phenotypes. (a) Results of the inter-group comparison of all parameters using the one-way ANOVA and Bonferroni post-hoc test. Differences between parameters with p<0.01 are highlighted in dark green. Differences with p<0.05 are indicated in light green. Only the parameters which could discriminate all three tumor models were used to generate confusion matrices. (b) Confusion matrices were generated to assess the capability of the parameters to correctly assign individual tumors to their according group. The numbers in the diagonal elements of the matrix represent correct classifications (highlighted in green), the remaining numbers indicate false assignments (highlighted in pink; see explanatory example in the upper left). Confusion matrices of the maximum of *distances to the closest vessel* and maximum of *distances to vessels with low velocities* reveal a correct classification in all cases (100%). For variance of *distances to the closest vessel*, 83% correct classification is achieved. (c) Although several parameters alone already allowed a correct assignment of all tumors, parameter combinations may be required when investigating more heterogeneous tumor populations. Therefore, a correlation matrix (Pearson’s correlation coefficient (r)) of all UBT parameters was generated to indicate the parameters providing complementary information. The highly discriminating distance parameters strongly correlated and thus, their combination may not be advantageous. However, the parameter *flow direction entropy* that distinguished two tumor models showed a low correlation with the distance parameters and could be selected as a potential candidate for a multi-parameter readout.

In order to determine the discriminatory power of the parameters at the basis of individual tumors, for each of these parameters the nearest neighbor classifier (NN) was applied in a leave-one-out-cross-validation, and confusion matrices were generated (Fig. 4b). It is clearly indicated that the maximum of *distances to the closest vessel* and maximum of *distances to vessels with low velocities* were best suitable for classifying three tumor models. With both parameters, a completely correct classification of all tumors was achieved (100%). With variance of *distances to the closest vessel*, 83% of the classifications were correct. However, all A431 tumors were classified correctly, and only one MLS tumor was wrongly assigned as an A549 tumor and one A549 tumor as a MLS tumor. Furthermore, with the mean of *distances to the closest vessel* and mean of *distances to vessels with low velocities*, 67% and 58% of the correct classification could be achieved, respectively (Fig. 4b).

In order to decide which parameters should be combined to correctly classify the tumors, it is important to investigate their interdependence. The lower the correlation between parameters that have high distinctive power, the higher is the probability that they will provide complementary information. For this purpose, a correlation matrix was generated. The superior distance parameters were all highly correlated and one single out of these parameters was sufficient to distinguish all tumors. Thus, a combination of parameters was not required. Nevertheless, it may become necessary to combine parameters during examinations of animals or patients with more heterogeneous tumors. In this context, a promising parameter is the *flow direction entropy*, since it can significantly distinguish two tumor groups and shows low correlations (r<0.5) with the distance parameters (Fig. 4c).

### Comparison of parameters derived from UBT and reference methods

To assess the robustness and the accuracy of UBT, we firstly compared the level of tumor vascularization (*rBV*) obtained by UBT to *rBV* values obtained by three other techniques, i.e. MIOT postprocessing, ex vivo micro computed tomography (μCT) and immunohistochemical (IHC) analysis of the tumor sections. Although *rBV* values did not differ significantly across the three tumor models, all modalities showed the same trend, classifying A431 tumors as the most vascularized ones, followed by MLS and A549 tumors (Fig 5a-e). However, at a quantitative scale *rBV* determined by MIOT revealed higher, μCT comparable and IHC slightly lower values than UBT (Fig. 5e).

**Fig. 5.**
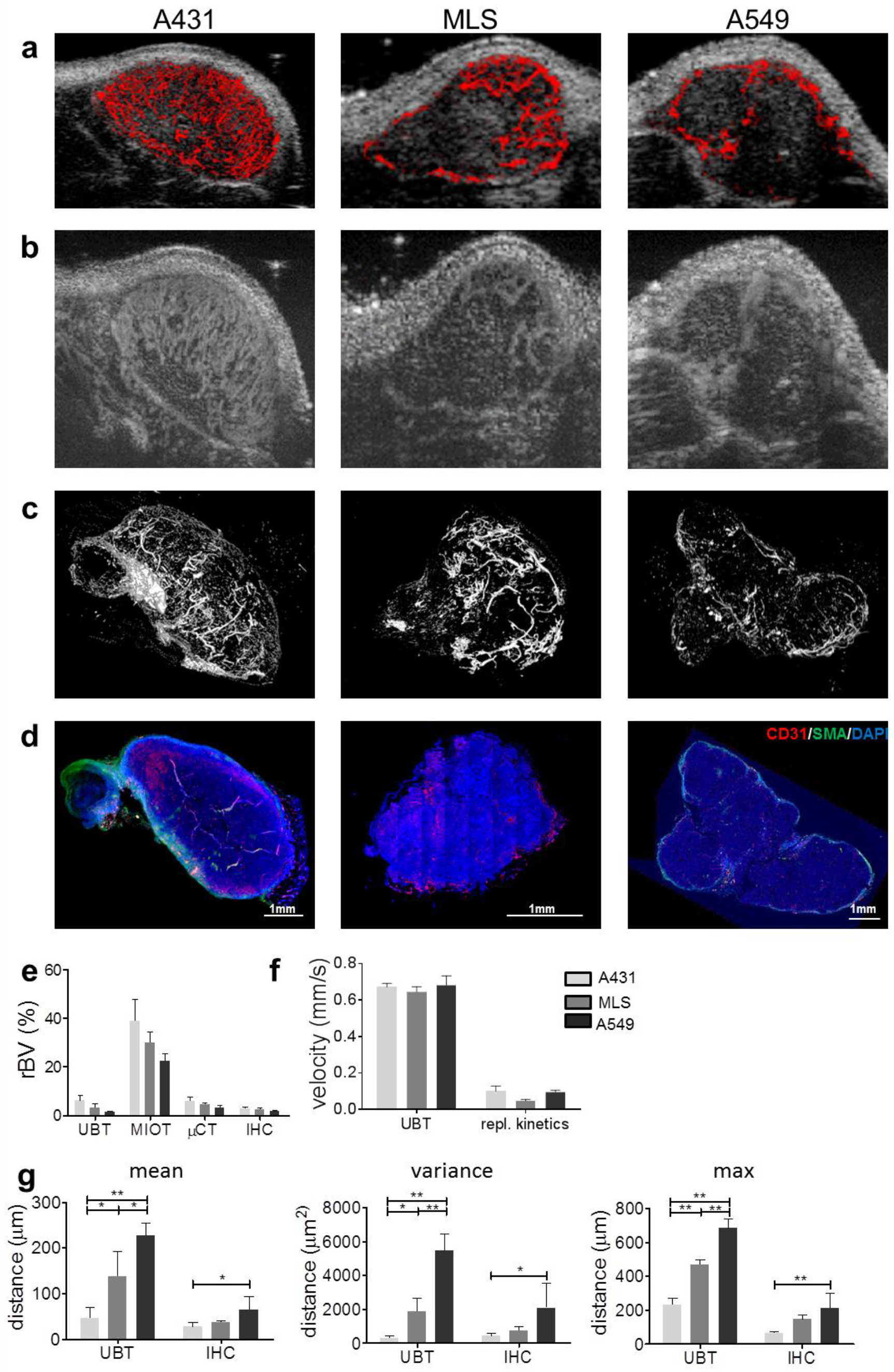
Comparison of UBT parameters with reference methods. *rBV* values in A431, MLS and A549 tumors (n=4 per tumor model) were obtained by UBT (a), Maximum Intensity Over Time (MIOT) US analysis (b), micro-computed tomography (μCT) (c), and immunohistochemistry (IHC)) (d). All methods show a similar trend, with A431 tumors having the highest and A549 tumors the lowest level of vascularization. While MIOT clearly overestimates the *rBV*, μCT and UBT provide comparable values, which are in line with the data from histology (e). Mean MB *velocity* values were either obtained from an exponential fit of a MB replenishment curve after a destructive US pulse from a ROI covering the entire tumor or from the UBT velocity maps (f). Both postprocessing procedures indicate that there are no significant differences in mean MB *velocities* between the tumor models. However, the absolute mean *velocity* values are significantly lower in the replenishment analysis than mean *velocities* calculated by UBT. Distance parameters determined by IHC analysis in the tumors with different vascular phenotypes are shown in (g). Mean, variance and maximum of the *distance to the closest vessel* determined by UBT had the same trend as their counterparts determined by IHC. While these parameters obtained by UBT analysis were significantly different across tumor models, by IHC analysis we could observe significant difference in A431 tumors compared to A549 (*=p<0.05, **=p<0.01) (data are presented as mean ± s.d. one-way ANOVA with Bonferroni post-hoc analysis).

In the next step, we compared mean *velocities* obtained by UBT with mean *velocities* calculated from replenishment kinetics. We found that both methods did not show differences in perfusion among the tumor models and provided values clearly below 1 mm/s. However, while UBT indicated mean *velocities* of approximately 0.8 mm/s, the values obtained by replenishment analysis were systematically lower (approximately 0.09 mm/s) (Fig. 5f).

Finally, the quantitative values (mean, variance and max) of *distances to the closest vessel* obtained by IHC und UBT presented with the same order, i.e. A431 had the smallest, A549 the largest and MLS intermediate values (Fig. 5g).

### Clinical proof of concept

In order to demonstrate that UBT can be performed using conventional US devices and clinically approved US contrast agents, three patients were investigated. The first patient was a 53-year old woman with an invasive “no special type (NST)” breast cancer examined after the third cycle of neoadjuvant chemotherapy with epirubicin and cyclophosphamide. The patient was scanned in B-mode using a 12 MHz transducer of the Toshiba Aplio 500 device (Toshiba Medical Systems GmbH, Otawara, Japan). Three milliliters of SonoVue (Bracco, Milan, Italy) were injected slowly over 6 min. Although the data were not acquired with a contrast-specific mode, UBT successfully captured a large vascular trunk with many feeding and draining vessels and thus identified the tumor areas that were still viable (Fig. 6a).

**Fig. 6.**
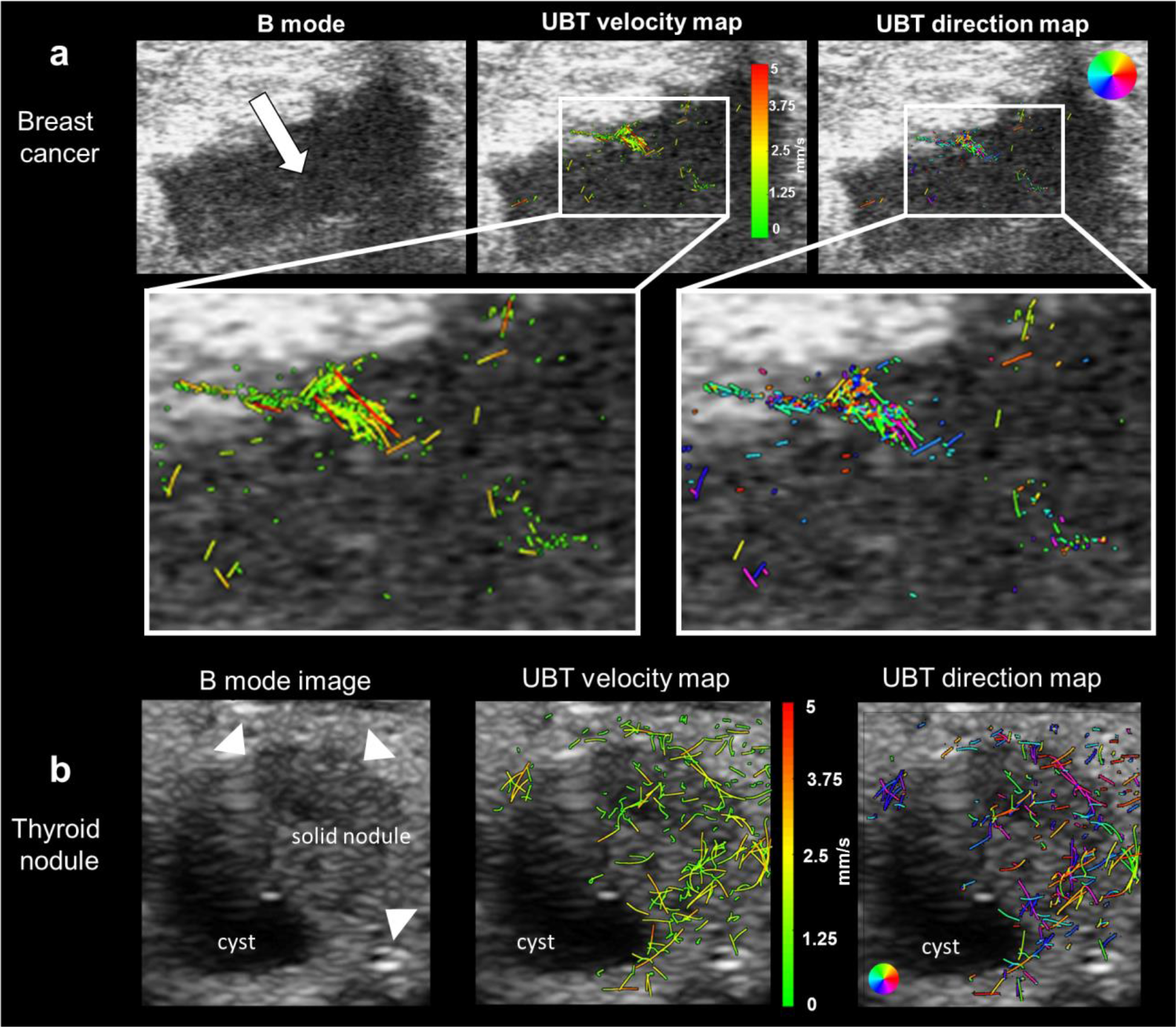
Preliminary clinical UBT images. (a) UBT images obtained in a patient with breast cancer. The upper row shows the B-mode image (left), the UBT velocity map (middle) and the UBT flow direction map of a breast cancer in patient using a conventional clinical US devices and phospholipid MB. The major part of the tumor shows a low vascularization, most possibly as the consequence of previous neoadjuvant chemotherapy. The bottom row presents magnifications of the most vascularized tumor part. Here a large vascular trunk comprising of the feeding and draining vessels with different velocities is displayed. (b) UBT images obtained in a patient with a cystic thyroid nodule. The nodule is presented with a solid component on the upper right part of the image and a cystic formation in the down-left angle. While no false tracks can be found in the cystic part, the vascular anatomy, vessel velocities, and flow directions are reliably depicted by UBT.

The second patient was a 30-year old woman with a thyroid nodule, scanned with a Zonare ZS3 (Zonare Medical Systems, Inc., Mountain View, CA) device using a 14 MHz transducer and a phase inversion/amplitude modulation contrast mode. After slow injection of SonoVue over 1 min, we could detect an avascular cystic formation filled with hypoechoic fluid in the lower left part of the nodule and a solid hypervascular part, where flow velocities and directions of individual MB could be visualized (Fig. 6b).

The third patient was a 56-year old woman with the Triple Negative Breast Cancer (TNBC) who received epirubicin/cyclophosphamide neoadjuvant chemotherapy every 3 weeks for 4 cycles. The patient was repeatedly imaged after the slow injection of 0.5 ml of SonoVue, before the initiation of the chemotherapy, after the first and after the second cycle with a 12 MHz transducer of the Toshiba Aplio 500 device. UBT super-resolution images nicely displayed the tumor vasculature and depicted the change in blood perfusion and flow direction over the course of treatment. At the baseline measurement, the vascularization was mainly located in the central part of the tumor without a dominant blood flow direction. Surprisingly, after the first chemotherapy cycle, we could observe apparent changes in the vascularization pattern. The tumor vascularization appeared much more homogeneous and strongly enhanced at the periphery. Additionally, rBV increased while the tumor size decreased from 25.6 cm^3^ to 3.0 cm^3^. After the second cycle of the chemotherapy the tumor size decreased remarkably to 0.8 cm^3^ and also its vascularization was substantially lower (Fig. 7). We hypothesize that tumor cell death induced by the first chemotherapy administration decreased the solid stress and/or interstitial fluid pressure ^19^, which then caused vascular decompression and thus, improved tumor perfusion and drug delivery in subsequent chemotherapy cycles.

**Fig. 7.**
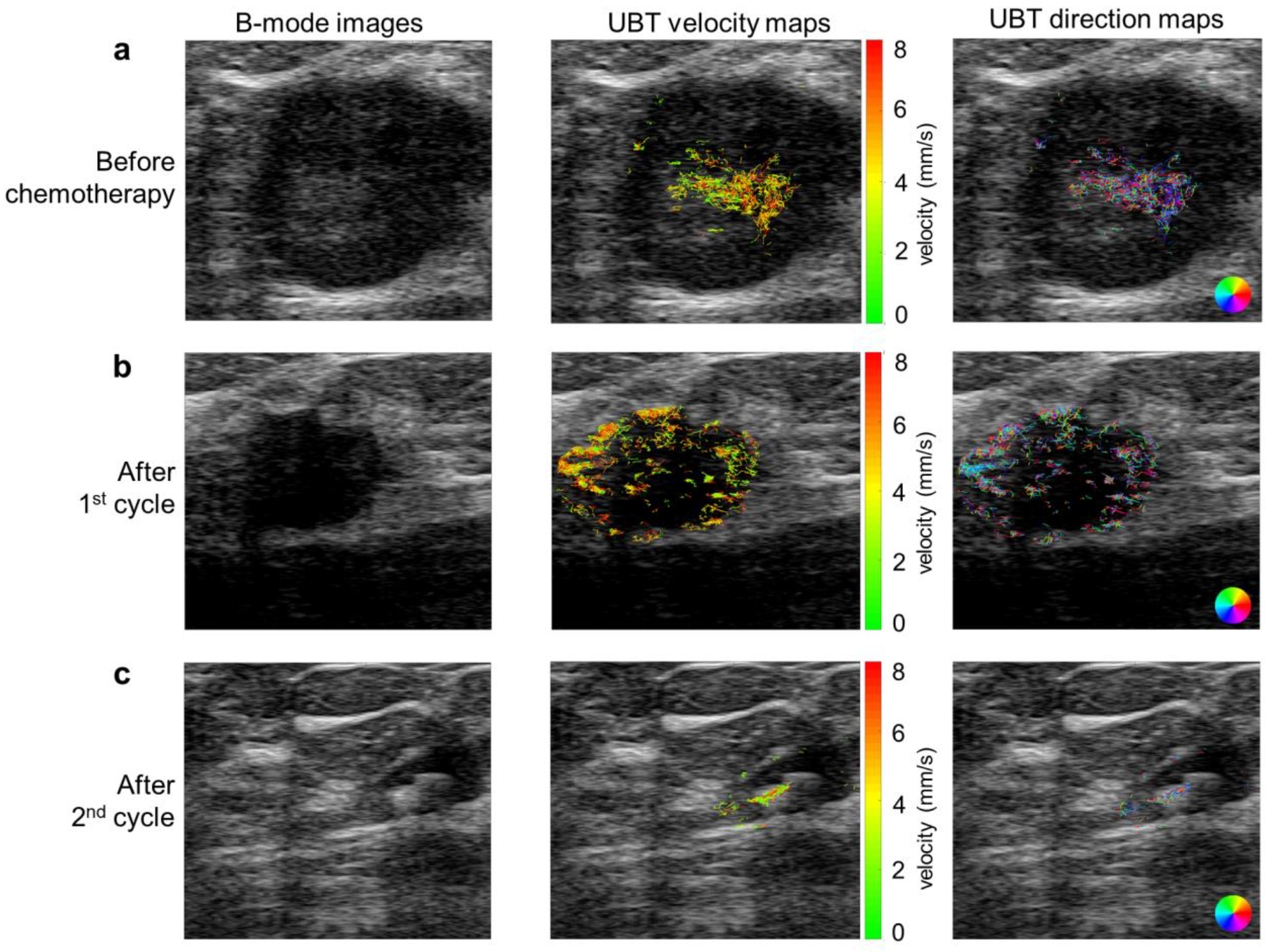
B-mode and UBT images of a triple negative breast carcinoma in a patient treated with neoadjuvant chemotherapy. CEUS measurements were performed with a conventional US device and phospholipid MB before (a), after the first (b) and after the second cycle (c) of chemotherapy. The first column shows B-mode images, the second column displays the UBT velocity maps and the third column indicates the UBT direction maps. At the baseline measurement, the tumor displayed a low vascularity and, only in its center, the vascular networks were depicted without showing any dominant direction (a). After the first cycle of treatment, the tumor size had decreased and vascularization appeared more homogeneous and more pronounced at the periphery (b). After the second cycle of treatment, the tumor had become very small and there was an obvious decline in the level of the vascularization (c).

## Discussion

In this study, we evaluated UBT for the structural and functional imaging of vascular features in murine and human tumors at super-resolution. We show that UBT opens new avenues for textural and functional tumor analysis at the individual vessel level and it provides novel classes of vascular parameters. In this context, reliable and robust quantification was achieved and different (vascular) phenotypes of tumors could be accurately discriminated. We postulate that this comprehensive and quantitative vascular characterization can be clinically highly valuable since the level of vascularization, microvessel density, and the distribution of vessels are often highly correlated with tumor invasiveness, aggressiveness, metastatic potential and prognosis of the disease ^20,21^, as already shown in different types of tumors e.g. brain tumors and melanoma ^22,23^.

When comparing UBT to reference techniques, we found that *rBV* values of tumors obtained by MIOT, μCT and IHC showed the same trend across the tumor models. At a quantitative scale *rBV* values obtained by UBT and μCT were very similar. However, as expected, MIOT overestimated the *rBV* since this technique counts every US voxel showing a positive MB signal as vessel, even if the vascular fraction within the respective voxel is small. In contrary, the somewhat lower *rBV* values obtained by IHC are explained by the fact that we preserved samples in formaldehyde, which is known to lead to tissue shrinkage ^24,25^.

Subsequently, mean *velocities* obtained by UBT and replenishment kinetics analysis were compared. In this context, it should be noted that in 25% of the cases, replenishment curves could not be fitted due to high noise levels in the US images, while all measurements were reliably postprocessed with UBT, which demonstrates its higher robustness. Nevertheless, both methods indicated that *velocity* values of the three tumor models were very similar. However, the values obtained by replenishment kinetics analysis were systematically lower than in UBT. Considering velocity values of mouse tumors reported in literature (1.1-1.5 mm/s) from multiphoton laser scanning microscopy ^26^, the quantitative numbers provided by UBT appear more realistic. In line with this, in previous publications the authors already reported that quantitative values obtained by replenishment kinetics analysis may not always be absolutely accurate ^26–28^. This may be explained by the fact that within a region of interest blood flow in the image plane cannot be detected and therefore remains unconsidered ^28^. Furthermore, the majority of replenishment analyses do not consider the influence of the beam elevation characteristics on the replenishment curve shape, which may also make the *velocity* values less accurate ^27^.

There is substantial need for further improvement of the UBT technique. In this context, our first patient measurements indicated several issues that, as long as unresolved, stand in the way of a broad clinical implementation. All measurements suffered from tissue motions that need to be compensated. While this is considerably easy for in-plane motion, out-of-plane movements cannot be corrected in 2D measurements, except when removing the non-matching slices, which leads to a loss of valuable data. Furthermore, the injection speed and concentration of MB need to be optimized. In case of the thyroid nodule, for example, the injection rate was too high, and individual MB could hardly be distinguished. Therefore, we used the CEUS sequences acquired in the early phase of the injection, which reduced the number of exploitable image frames. Consequently, some vessel trees might not have been completely reconstructed. Thus, CEUS scans for UBT should be performed under slow MB injection and will require much lower MB doses than for conventional methods, which in turn, however, may help to decrease potential side effects and concerns that have been raised, e.g. for using CEUS in patients with instable cardiopulmonary conditions and pulmonary hypertension ^29^. Furthermore, our data suggest that in clinical settings, contrast-specific modes should be applied to acquire US data for UBT, since the low signal-to-noise ratio of the B-mode, as used for the first case (breast cancer), made the MB detection more difficult than in the contrast mode scan used for the second (thyroid nodule) and third patient (neoadjuvant breast cancer therapy). Additionally, we observed relatively high velocities in the thyroid nodule measurements and therefore, a frame rate of about 17 Hz was close to the bottom threshold level for achieving reliable MB tracks. Another translational challenge represents the slice thickness of the image plane, which is larger in the clinical US scanners than in preclinical US systems, and therefore, significant overlays of vessels are expected. This makes it difficult to correctly connect the tracks of different MB. To overcome this problem, Lin et al. detected positions of the MB at super-resolution in 3D with a stepwise motorized motion stage, a high frame-rate system and a long acquisition time ^30^. Although they generated 3D super-resolution images of the vasculature, the individual MB were not tracked over time, thus the quantitative information about the hemodynamics was not obtained. We believe that the use of matrix transducers ^31^ for 3D UBT measurements may represent the ideal way to reconstruct and quantify the vascular network more completely and accurately.

In summary, our results demonstrate that UBT is a robust and reliable method that can be applied to data of commercial US systems. UBT can depict and accurately quantify important characteristics of the tumor vascularization at the individual vessel level and can generate new classes of vascular biomarkers that show superior performance over other CEUS methods in discriminating tumors. By providing super-resolution images of tissue vascularization, UBT offers new opportunities for a robust pattern analysis in US imaging and, in our opinion, has the potential to become an indispensable tool in tumor diagnosis and therapy monitoring. Moreover, UBT may not only be applied in oncology but may also be relevant for other indications, e.g. for characterizing inflamed tissues (e.g. inflammatory bowel disease), organ fibrosis (e.g. in liver and kidney), and immunological disorders (e.g. rheumatological disorders and organ rejection after transplantation). Furthermore, it may be applied to monitor vascular remodeling in the cardiovascular field (e.g. revascularization in ischemic tissues) and to assess antiangiogenic treatment effects e.g. in retinopathies. Based on the presented data, we are confident that, after further refinements, UBT will experience a rapid translation into clinical practice.

## Materials and Methods

### Study design

The objective of this study was to establish UBT as a CEUS postprocessing method for distinguishing tumors with different vascular phenotypes at super-resolution, as well as to provide proof of principle for applying UBT on clinical data. The study consisted of three major parts. In the first part, the ability of UBT to distinguish tumor models with different vascular phenotypes was assessed ^32^. For this purpose, A431, MLS and A549 tumor xenografts were induced in female immunodeficient CD1-nude mice (n=4 mice per tumor model). CEUS imaging was performed, and various parameters were extracted that describe morphological and functional vascular characteristics. Statistical tests were applied to investigate the diagnostic potential of these parameters for discriminating three tumor models and to find their ideal combination. In the second part of the study, we evaluated the robustness of UBT and the accuracy of the outcome parameters using available reference techniques. For that purpose, CEUS cine loops were analyzed with MIOT and replenishment kinetics to assess *rBV* values and mean *velocities*, respectively. For further validation of the *rBV* values, high resolution μCT scans of Microfil perfused tumors and histological analyses of tumor sections were evaluated (n=4 mice per tumor model). In the third part of the study, we applied our technology on CEUS data from three patients, in order to demonstrate its translational potential.

### Study approval

All animal experiments were approved by the local and governmental committee on animal care. The patients' examinations with CEUS were performed as part of the clinical routine diagnostic procedure. All patients gave their informed consent to retrospectively extract the relevant US images from the stored routine data and use them for the image postprocessing.

### Cell culture

The human cancer cell lines, A431 (epidermoid carcinoma), MLS (ovarian carcinoma) and A549 (non-small cell lung carcinoma) were obtained from American Type Culture Collection (Manassas, VA). A431 and MLS tumor cells were maintained in Roswell Park Memorial Institute 1640 medium (RPMI) and α-Minimum Essential Medium (α-MEM), respectively, and A549 cells were cultivated in Dulbecco’s Modified Eagle Medium (DMEM). All media (Life Technologies, Darmstadt, Germany) were supplemented with 10% fetal bovine serum and 1% Penicillin/Streptomycin (Gibco, Invitrogen, Germany). Cells were incubated at 37°C in 5% CO2 and passaged at 80-90% confluence.

### Xenograft tumor models

The mice were housed in groups of four per cage under specific pathogen-free conditions with a 12h light and dark cycle in a temperature- and humidity-controlled environment (according to FELASA-guidelines). Water and standard pellets for laboratory mice (Sniff GmbH, Soest, Germany) were offered ad libitum. Human tumor xenografts were induced in 8-weeks old female immunodeficient CD1-nude mice (Crl:CD1-Foxn1nu, Charles River, Sulzfeld, Germany) (n=4 mice per tumor model). For this purpose, 4×10^6^ A431, MLS or A549 tumor cells were injected subcutaneously into the right flank. When tumors reached a size of approximately 5×5 mm, CEUS imaging experiments were performed. Prior to experiments, animals were anesthetized by inhalation of 2% isoflurane in oxygen.

### Contrast-enhanced ultrasound imaging

Hard-shell polybutylcyanoacrylate (PBCA) MB were used as US contrast agent to assess the potential of UBT for tumor characterization. PBCA-MB were freshly synthetized as described before ^33^. For animal experiments, the PBCA-MB suspension was diluted in sterile sodium chloride to a concentration of 2×108 MB/ml. Each mouse was injected with a 50 μL bolus containing 1×10^7^ PBCA-MB over approximately 3 seconds, followed by a 20 μl saline flush, into the lateral tail vein.

### Image acquisition

For the US measurements, a Vevo 2100 system equipped with the MS-550D probe (FUJIFILM Visualsonics, Toronto, ON, Canada) was used. The linear array exhibited a center frequency of 40 MHz and a bandwidth from 22 MHz to 55 MHz. The maximum image depth was 15 mm.

The correct placement of the probe on the tumors was controlled prior to the measurements using real-time B-mode imaging. For the measurements, image series were recorded during the destruction-replenishment sequence at a frame rate of 50 frames/s. The images were acquired in digital raw radio frequency (RF) mode to receive uncompressed IQ data. The gain was set to 22 dB and the transmit power to 2% to minimize MB destruction. For each tumor, the number of processed frames was limited to 2000, which was equivalent to 40 s measurement time.

### Image processing

After the US measurements, the tumor border was outlined manually and the further processing steps were carried out inside the segmented area.

In a first step, a rigid motion estimation and compensation were carried out. For this, the B-mode images were interpolated on a finer grid (4-fold) to increase the accuracy of the motion compensation. The motion profiles over time typically exhibited periods of small movements disturbed by spikes of large movements due to breathing. These frames of large movements were excluded because they typically included also out-of-plane movements.

To detect single echoes of individual MB, the B-Mode images were separated into static background and moving foreground images. The rolling background was calculated applying a temporal rank filter of rank 3 over ±10 frames around the actual frame. The foreground was computed by subtracting the background from the original frames. After applying an adaptive threshold to the foreground images, the MB were localized by calculating the intensity weighted centroid for each MB. Choosing a too low threshold leads to false detections due to noise in the image, choosing a too high threshold leads to missed detections. Therefore, for each measurement set, the threshold was adapted to result in no detections immediately after the destruction event in the destruction replenishment sequence.

### Tracking of microbubbles

To reconstruct the vasculature, the detected MB must be tracked over several frames. As discussed in ^16^, usually this is either achieved by a nearest-neighbor tracking for very high frame rates in the kHz range, which need non-standard ultrafast US scanners, or by very low MB concentrations and long observation times up to several minutes.

Here, we solved the tracking challenge that occurs when using clinically recommended MB concentrations and standard US systems, by using a novel, more robust UBT method. This was necessary, since tracking by using the nearest MB in the next frame to continue a track is prone to failure under the given imaging conditions: Due to the elevational width of the imaging slice, an apparent crossing of capillaries is expected when capillaries of different directions are running in different planes within a slice. Furthermore, the number of MB can be high (up to 100 MB per frame) for a small tumor area (e.g. ~ 14 mm^2^) leading to a high MB density of > 7 MB per mm^2^. Thus, considering the applicable frame rate and the expected flow velocities, the closest MB will often not be the correct one to continue the track. Therefore, a more robust Markov Chain Monte Carlo Data Association (MCMCDA) algorithm was applied to the detected MB positions to track the MB over several frames ^34^. This algorithm associates detected positions based on a probabilistic optimization considering a motion model. A detailed description of the algorithm can be found in ^34^ but the main concepts are briefly described: Bubble positions can be associated in different ways to the tracks, as illustrated in Fig. 1A by two different track associations *ω*_1_ and *ω*_2_. The algorithm evaluates the a-posteriori probability *P*(*ω*|*Y*) of an association ω under the given measured positions Y. By Bayes’ rule, this probability is proportional to

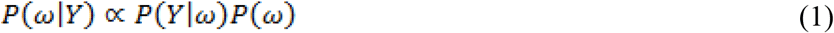

In this, *P*(*ω*|*Y*) is the likelihood of the measured position Y under a given track association as determined from a linear motion model. The a priori probability of a track association, without the knowledge of the measurement data, is is *P*(*ω*), which is calculated from the assumed probabilities of false detections, missing detections, track starts, and track ends. These parameters of the algorithm are chosen as described in ^34^.

Trying all possible track combinations and finding the association *ω*_max_ that maximizes the a posteriori probability: *ω_max_* = arg max_*ω*∊Ω_ *P*(*ω*|*Y*) is, however, an intractable combinatorial problem. Thus, in a Monte Carlo approach a Markov chain was used to randomly draw associations with this probability distribution. By this, associations with high probability are drawn more often and the best association will be kept. For example, in Fig. 1a, the track association *ω*_1_ had a higher probability compared to the nearest neighbor association represented as *ω*_2_, because the position prediction of the motion model resulted in higher likelihoods P(Y|ω) of the measured positions under the assumption of track association ω, which also led to a higher a posteriori probability *P*(*ω*|*Y*).

The tracking algorithm yielded not only the trajectories but also the velocities and directions of the moving MB.

### Definition and extraction of parameters

For further evaluations, images of MB tracks, velocities, and flow direction were reconstructed with a pixel size of 5 × 5 μm^2^ for each tumor. Even though we were able to achieve a higher resolution with the 40 MHz transducer, we used these settings due to the MB’s size (3μm), the limited resolution in the graphical illustrations and to restrict the data amount.

Microbubble track images were generated using Bresenham’s line algorithm to connect the MB positions of the estimated tracks. The MB track image was a binary image indicating the pixels which were passed by the MB. In the flow velocity map and the flow direction map, the corresponding quantity was assigned to the pixels along the track.

For the evaluation of the parameters, each tumor was divided into two regions: a rim of 0.5 mm thickness and the core which was the remaining area when excluding the rim from the whole tumor area. For the characterization of the tumor vasculature, we used only the core region to exclude the large feeding vessels in the rim.

From the MB track image the *rBV* was derived as the ratio of the area covered by the tracks to the respective total area, which was expected to be proportional to the *rBV*.

Additionally, from the MB track image a track distance map was generated applying the Euclidean distance transform (*bwdist* function, Matlab, MathWorks, Natick, MA, USA). For each pixel, the track distance map provided the shortest distance to the next vessel. For each track distance map, mean, variance, maximum, and median of the *distances to the closest vessel* were calculated for the respective areas. Small distances are characteristic for a fine meshwork of vessels. The larger the maximum distance, the larger are the non-perfused areas.

From the flow velocity map the statistics of the velocities were derived. Again, mean, variance, maximum and median were calculated.

Since we were interested in the structure of the vasculature, we divided the vessels into two groups of high and low flow velocities, respectively, and calculated the distance parameters separately for the resulting two groups. We defined the mean value of the median velocities of the tumors as the threshold between high and low flow velocities (0.7 mm/s) and calculated the mean, variance, maximum, and median of the *distances to vessels with low and high velocities*.

We were also interested in parameterizing the directions of MB flow. To characterize a locally ordered flow with predominant directions in contrast to locally chaotic flow directions, we defined sub-regions of 25 μm × 25 μm and calculated the *flow direction entropy* of the vessels from the flow direction maps. Predominant directions within sub-regions will result in low entropy values. Local entropy values are averaged over the tumor core area.

### Statistical analysis

Data are presented as mean ± standard deviation (s.d). The one-way ANOVA and Bonferroni post-hoc test were applied to evaluate differences between groups considering a p-value of <0.05 to be significant. All analyses were performed using GraphPad Prism 5.0 (GraphPad Software, San Diego, CA).

### Confusion and correlation matrices

For all parameters extracted by UBT, one-way ANOVA and Bonferroni post-hoc tests were applied. Parameters that distinguish all three tumor models were used for the further analyses. Then, for each parameter the nearest neighbor classifier (NN) was applied in an exhaustive leave-one-out-cross-validation. The results are presented in confusion matrices, which plot the actual classes versus the classes predicted by the classification. The numbers in the diagonal elements of the matrix represent the correct classifications; the remaining numbers indicate the false assignments. The correct classification is expressed in percentage.

Additionally, a correlation matrix of all parameters obtained by UBT was generated to depict the pairwise dependencies among them measured by the Pearson’s correlation coefficient (r).

Confusion and correlation matrices were generated by Matlab, MathWorks, Natick, MA, USA.

### Reference methods

We firstly validated *rBV* obtained by UBT by *rBV* obtained by MIOT analysis, ex vivo μCT and histological evaluation of the tumor sections. For generation of the MIOT images, the highest amplitude values of each pixel were preserved throughout the recorded B-mode image sequence after MB injection. Then, from each CEUS image a corresponding background image, containing the median pixel-wise value of all B-mode images, was subtracted to reveal the vascular network. To assess the *rBV* from MIOT images, a threshold-based segmentation of the vessels was performed and *rBV* was calculated as the fraction of vessels in the tumor area using the Imalytics Preclinical software (Gremse-IT, Aachen, Germany) ^35^.

Next, in terminal experiments, mice were perfused intracardially with the silicone rubber radiopaque compound Microfil^®^ (FlowTech, Carver, MA), which polymerizes in blood vessels within 20 min ^32^. After Microfil perfusion, tumors were excised, preserved in 4% formalin and scanned in the high-resolution desktop X-ray micro-CT system SkyScan 1172 with a Hamamatsu 10 Mp camera (pixel size 11.66 μm) (SkyScan, Kontich, Belgium). Tumors were scanned around the vertical axis with rotation steps of 0.3° at 59 kV and a source current of 167 μA. 640 projections (2096×4000 pixels) were acquired during 2.5-3 hours per tumor. After threshold-based segmentation, *rBV* was determined as fraction of Microfil-perfused vessels of total tumor volume using the Imalytics Preclinical software (Gremse-IT, Aachen, Germany) ^35^.

Finally, tumors were embedded in paraffin and cut into 5 μm thick sections. Immunostaining of endothelial cells was performed with a rat anti-mouse CD31 primary antibody (BD Biosciences, Heidelberg, Germany), followed by a donkey anti-rat Cy-3-labeled secondary antibody (Dianova, Hamburg, Germany). Smooth muscle cells and pericytes were labelled using a biotinylated anti-α-smooth muscle actin (α-SMA) primary antibody (Progen, Heidelberg, Germany) and streptavidin-Cy-3 (Dianova, Hamburg, Germany). Nuclei were counterstained with 4, 6-diamidino-2-phenylindole (DAPI; Invitrogen, Karlsruhe, Germany). Fluorescent micrographs were obtained with an Axio Imager M2 light microscope and AxioCamMRm revision 3 high-resolution camera (Carl Zeiss Microimaging, Göttingen, Germany). For each tumor a whole histopathological section was analyzed. The vessel area fraction, referring to *rBV*, was calculated by semi-automated detection and filling of the lumen of CD31-positive structures and dividing the resulting area by the total tumor area. Furthermore, the vessel maturity index was determined by calculating the percentage of SMA positive vessels per total number of vessels. All IHC analyses were performed using the AxioVision Rel 4.8 software (Carl Zeiss Microimaging).

Secondly, mean *velocities* obtained by UBT were compared to mean MB *velocities* calculated by destruction replenishment analysis, which is a clinically established US method. The corresponding algorithm was implemented in a custom program (Matlab, R2015a, MathWorks, Natick, USA). Cine loops were acquired over 20 seconds (50 frames per second, 1000 frames in total) with a 40 MHz transducer. After recording the MB bolus injection phase, a destructive pulse was applied for 1 second to destroy all MB in the imaged tumor slice. Then, the replenishment of circulating MB was recorded over approximately 40 seconds. *Velocities* were determined by fitting the slope of the replenishment curve as described by Wei et al ^12^.

Finally, we validated the *distances to the closest vessel* obtained by UBT to their counterparts obtained by IHC analysis. The *distances to the closest vessel* were calculated manually from 5 micrographs of each histopathological section, as the shortest distance from one CD31 positive to the next CD31 positive vessel wall. Then we calculated mean, variance and maximum values. IHC analyses were performed using the AxioVision Rel 4.8 software (Carl Zeiss Microimaging).

### Patient examinations

The patient with the thyroid nodule had been examined at the Department of Radiology of the German Cancer Research Center in Heidelberg, Germany, and the patients with breast cancer were scanned at the Department of Gynecology and Obstetrics at the University Medical Center of the RWTH University in Aachen, Germany. All further details about the scanning procedure are specified in the results.

### Data availability

The data that support the findings of this study are available from the corresponding authors on request.

## Acknowledgments

The authors thank M. Weiler for performing μCT measurements and S. von Stillfried for providing paraffin embedded samples. This study is supported by Deutsche Forschungsgemeinschaft DFG (KI1072/11-1).

## Author contributions

F.K. and G.S. planned the study, supervised the experiments and the data analysis, and revised the manuscript. T.O. and A.R. performed the experiments. S.D., D.A., M.P., and G.S. established the algorithm and preformed the postprocessing analyses. E.S. and S.Del. provided the clinical data. B.T. and A.R. assisted with the data analyses and reviewed the manuscript. T.O., S.D., and M.P. prepared the figures. T.O., and S.D. drafted the manuscript. **Competing interests:** The authors declare that they have no competing interests.

## References and Notes

1 Lindner, J. R. Microbubbles in medical imaging: current applications and future directions. Nat Rev Drug Discov 3, 527–532, doi:10.1038/nrd1417 (2004).

2 Padhani, A. R., Harvey, C. J. & Cosgrove, D. O. Angiogenesis imaging in the management of prostate cancer. Nat Clin Pract Urol 2, 596–607, doi:10.1038/ncpuro0356 (2005).

3 Hamer, O. W., Schlottmann, K., Sirlin, C. B. & Feuerbach, S. Technology insight: advances in liver imaging. Nat Clin Pract Gastroenterol Hepatol 4, 215–228, doi:10.1038/ncpgasthep0766 (2007).

4 Dimastromatteo, J., Brentnall, T. & Kelly, K. A. Imaging in pancreatic disease. Nat Rev Gastroenterol Hepatol 14, 97–109, doi:10.1038/nrgastro.2016.144 (2017).

5 Birner, P. et al. Vascular patterns in glioblastoma influence clinical outcome and associate with variable expression of angiogenic proteins: evidence for distinct angiogenic subtypes. Brain Pathol 13, 133–143 (2003).

6 Tolaney, S. M. et al. Role of vascular density and normalization in response to neoadjuvant bevacizumab and chemotherapy in breast cancer patients. Proc Natl Acad Sci U S A 112, 14325–14330, doi:10.1073/pnas.1518808112 (2015).

7 Tadeo, I. et al. Vascular patterns provide therapeutic targets in aggressive neuroblastic tumors. Oncotarget 7, 19935–19947, doi:10.18632/oncotarget.7661 (2016).

8 Jia, W. R. et al. Three-dimensional Contrast-enhanced Ultrasound in Response Assessment for Breast Cancer: A Comparison with Dynamic Contrast-enhanced Magnetic Resonance Imaging and Pathology. Sci Rep 6, 33832, doi:10.1038/srep33832 (2016).

9 Jain, R. K. et al. Biomarkers of response and resistance to antiangiogenic therapy. Nat Rev Clin Oncol 6, 327–338, doi:10.1038/nrclinonc.2009.63 (2009).

10 Mirnezami, R., Nicholson, J. & Darzi, A. Preparing for precision medicine. N Engl J Med 366, 489–491, doi:10.1056/NEJMp1114866 (2012).

11 Pysz, M. A. et al. Assessment and monitoring tumor vascularity with contrast-enhanced ultrasound maximum intensity persistence imaging. Invest Radiol 46, 187–195, doi:10.1097/RLI.0b013e3181f9202d (2011).

12 Wei, K. et al. Quantification of myocardial blood flow with ultrasound-induced destruction of microbubbles administered as a constant venous infusion. Circulation 97, 473–483 (1998).

13 Konerding, M. A., Miodonski, A. J. & Lametschwandtner, A. Microvascular corrosion casting in the study of tumor vascularity: a review. Scanning Microsc 9, 1233–1243; discussion 1243-1234 (1995).

14 Siepmann, M., Schmitz, G., Bzyl, J., Palmowski, M. & Kiessling, F. in 2011 IEEE International Ultrasonics Symposium. 1906–1909.

15 Theek, B. et al. Automated Generation of Reliable Blood Velocity Parameter Maps from Contrast-Enhanced Ultrasound Data. Contrast Media &#x26; Molecular Imaging 2017, 8, doi:10.1155/2017/2098324 (2017).

16 Errico, C. et al. Ultrafast ultrasound localization microscopy for deep super-resolution vascular imaging. Nature 527, 499–502, doi:10.1038/nature16066 (2015).

17 Christensen-Jeffries, K., Browning, R. J., Tang, M. X., Dunsby, C. & Eckersley, R. J. In vivo acoustic super-resolution and super-resolved velocity mapping using microbubbles. IEEE Trans Med Imaging 34, 433–440, doi:10.1109/TMI.2014.2359650 (2015).

18 Jain, R. K., Martin, J. D. & Stylianopoulos, T. The role of mechanical forces in tumor growth and therapy. Annu Rev Biomed Eng 16, 321–346, doi:10.1146/annurev-bioeng-071813-105259 (2014).

19 Stylianopoulos, T. & Jain, R. K. Combining two strategies to improve perfusion and drug delivery in solid tumors. Proc Natl Acad Sci U S A 110, 18632–18637, doi:10.1073/pnas.1318415110 (2013).

20 Ruoslahti, E. Specialization of tumour vasculature. Nat Rev Cancer 2, 83–90, doi:10.1038/nrc724 (2002).

21 Carmeliet, P. & Jain, R. K. Molecular mechanisms and clinical applications of angiogenesis. Nature 473, 298–307, doi:10.1038/nature10144 (2011).

22 Kashani-Sabet, M., Sagebiel, R. W., Ferreira, C. M., Nosrati, M. & Miller, J. R., 3rd. Tumor vascularity in the prognostic assessment of primary cutaneous melanoma. J Clin Oncol 20, 1826–1831, doi:10.1200/JCO.2002.07.082 (2002).

23 Chow, K. L. et al. Prognostic factors in recurrent glioblastoma multiforme and anaplastic astrocytoma treated with selective intra-arterial chemotherapy. AJNR Am J Neuroradiol 21, 471–478 (2000).

24 Boonstra, H., Oosterhuis, J. W., Oosterhuis, A. M. & Fleuren, G. J. Cervical tissue shrinkage by formaldehyde fixation, paraffin wax embedding, section cutting and mounting. Virchows Arch A Pathol Anat Histopathol 402, 195–201 (1983).

25 Hsu, P. K. et al. Effect of formalin fixation on tumor size determination in stage I non-small cell lung cancer. Ann Thorac Surg 84, 1825–1829, doi:10.1016/j.athoracsur.2007.07.016 (2007).

26 Kamoun, W. S. et al. Simultaneous measurement of RBC velocity, flux, hematocrit and shear rate in vascular networks. Nat Methods 7, 655–660, doi:10.1038/nmeth.1475 (2010).

27 Hudson J.M., K. R., Burns P.N. Quantification of flow using ultrasound and microbubbles: a disruption replenishment model based on physical principles. Ultrasound Med Biol 35, 2007–2020 (2010).

28 Krix, M. et al. A multivessel model describing replenishment kinetics of ultrasound contrast agent for quantification of tissue perfusion. Ultrasound Med Biol 29, 1421–1430 (2003).

29 Claudon, M. et al. Guidelines and good clinical practice recommendations for Contrast Enhanced Ultrasound (CEUS) in the liver – update 2012: A WFUMB-EFSUMB initiative in cooperation with representatives of AFSUMB, AIUM, ASUM, FLAUS and ICUS. Ultrasound Med Biol 39, 187–210, doi:10.1016/j.ultrasmedbio.2012.09.002 (2013).

30 Lin, F. et al. 3-D Ultrasound Localization Microscopy for Identifying Microvascular Morphology Features of Tumor Angiogenesis at a Resolution Beyond the Diffraction Limit of Conventional Ultrasound. Theranostics 7, 196–204, doi:10.7150/thno.16899 (2017).

31 Wang, H., Lutz, A. M., Hristov, D., Tian, L. & Willmann, J. K. Intra-Animal Comparison between Three-dimensional Molecularly Targeted US and Three-dimensional Dynamic Contrast-enhanced US for Early Antiangiogenic Treatment Assessment in Colon Cancer. Radiology 282, 443–452, doi:10.1148/radiol.2016160032 (2017).

32 Ehling, J. et al. Micro-CT imaging of tumor angiogenesis: quantitative measures describing micromorphology and vascularization. Am J Pathol 184, 431–441, doi:10.1016/j.ajpath.2013.10.014 (2014).

33 Fokong, S. et al. Advanced characterization and refinement of poly N-butyl cyanoacrylate microbubbles for ultrasound imaging. Ultrasound Med Biol 37, 1622–1634, doi:10.1016/j.ultrasmedbio.2011.07.001 (2011).

34 Ackermann D., S. G. Detection and Tracking of Multiple Microbubbles in Ultrasound B-Mode Images. IEEE Trans Ultrason Ferroelectr Freq Control 63, 72–82 (2016).

35 Gremse, F. et al. Imalytics Preclinical: Interactive Analysis of Biomedical Volume Data. Theranostics 6, 328–341, doi:10.7150/thno.13624 (2016).

